# Association Between Rare Copy Number Variation and Response to Social Skills Training in Autism Spectrum Disorder

**DOI:** 10.1101/380147

**Authors:** Kristiina Tammimies, Danyang Li, Ielyzaveta Rabkina, Sofia Stamouli, Martin Becker, Veronika Nicolaou, Steve Berggren, Christina Coco, Torbjörn Falkmer, Ulf Jonsson, Nora Choque-Olsson, Sven Bölte

## Abstract

Challenges in social communication and interaction are core symptoms in autism spectrum disorder (ASD) for which social skills group training (SSGT) is a commonly used intervention. SSGT has shown modest but heterogeneous effects in clinical trials, and therefore identification of effect moderators could enable more precise intervention decisions. One of the major genetic risk factors in ASD are rare copy number variation (CNV). However, limited information exists whether rare CNVs profiles can be used to aid in intervention decisions. Therefore, we conducted the first study to date analyzing rare CNVs as genetic moderators in the outcome of SSGT in ASD. For this, we analyzed rare genic CNV carrier status of 207 children of which 105 received SSGT and 102 standard care as part of a recent randomized clinical trial for 12-weeks SSGT. We used mixed linear models to assess the association of being a CNV carrier, grouped by the effect and size of the CNVs and the primary response to SSGT, the parent-report Social Responsiveness Scale (SRS) measured at post-intervention and 3-months follow-up. Additionally, we analyzed the secondary outcome assessments included parent-rated adaptive behaviors (ABAS-II) and trainer-rated clinical global impression (CGI). We show that being a carrier of any size rare genic CNV did not impact on the SSGT outcome. However, when stratifying the groups by size of the CNVs, we identified that carriers of large CNVs (>500 kb) showed inferior SRS outcomes at post-intervention (β = 15.35, 95% CI 2.86-27.84, *P*=0.017) and follow-up (β = 14.19, 95% CI 1.68-26.70, *P*=0.028). Similar results were shown for the parent-rated secondary outcome. In contrast, the carriers of small CNVs had better outcome at post-intervention (β = −1.20, 95 *%* CI - 2.0 - −0.4 *P* = 0.003) but not at follow-up for the trainer-rated secondary outcome CGI. These results remained when we tested the specificity of the effect by including the standard care group and adjusting for IQ levels. While our study suggests that being a carrier of any size rare genic CNV did not impact the outcome, it provides preliminary evidence that carriers of high-risk CNVs might not benefit on SSGT as much as non-carriers. Our results indicate that genetic information eventually might help guide personalized intervention planning in ASD. We additionally highlight that more research is needed to understand the intervention needs of autistic individuals with specified molecular alterations.

## Introduction

Autism spectrum disorder (ASD) is a neurodevelopmental condition with a prevalence of 1-2%.^1,2^ A multitude of alterations in social communication and interaction are among the core challenges in ASD.^3^ The clinical presentation is heterogeneous, and the condition often co-occurs with other complications, such as attention-deficit hyperactivity disorder (ADHD), anxiety and depression, intellectual disability, and variety of somatic disorders.^4,5^ Reflecting the nature of the spectrum of clinical manifestations, the causes of ASD are diverse. Genetic factors, both common and rare play a significant part in the etiology of the disorder with complex interplay with each other and environmental factors.^6,7^

There is a scarcity of evidence-based interventions for ASD.^8^ A commonly used intervention to ameliorate social communication difficulties and their negative impact on adaptive outcomes in the normative intellectual range of ASD is social skills group training (SSGT). SSGT is based on a combination of behavioral modification and socially instructive techniques applied in cohesive and safe peer group settings. The accumulated body of randomized controlled trials (RCTs) on SSGT suggest that the average effect is modest.^9-11^ However, the overall heterogeneity of the outcome suggests that subgroup analyses might help identify moderators and mediators. The largest RCT on SSGT to date is the KONTAKT program^12^ led by our center, found that the effect on social skills of a 12-week training was restricted to adolescents and females.^10^ Further, a previous uncontrolled trial of SSGT KONTAKT found language capacities to be positively correlated with outcome, while ASD severity negatively impacted the outcome.^13^ Another previous SSGT trial did not find the outcome to be associated with ADHD, level of social anxiety, Theory-of-Mind, or desire for social interaction.^14^ However, the sample size was insufficient for informative subgroup analyses.

Genetic factors have not yet been studied as potential sources of heterogeneity in SSGT outcomes. The most convincing genetic risk markers in ASD are rare copy number variants (CNVs), deletions and duplications of genetic material. Rare CNVs confers up to 20-fold increase risk for ASD.^15^ The most frequent CNVs underlying ASD are 16p13.2 microduplication (MIM: 613458), 15q11-q13 duplication (MIM: 608636), 1q21.1 deletion/duplication (MIM: 612474/612475) and 16p11.2 deletion (MIM:611913) syndromes.^16^ Therefore, chromosomal microarray analysis (CMA) is recommended as the first-tier genetic test to identify CNVs in ASD with a molecular diagnostic yield up to 25%.^17-19^ In addition to providing information about the genetic causes and recurrence risks for family members, CMA has been reported to result in individual medical intervention recommendations such as referral to specialists, surveillance protocols, and family investigations.^20,21^

An unanswered question of high relevance for clinical practice is whether CMA results can guide the selection and prioritization of interventions in ASD. To address this question, we performed analyzes of the potential moderating effect of CNVs on the outcome of a cohort following a 12-weeks training of SSGT KONTAKT. To increase the probability of clinically useful results, we focused on rare genic CNVs, including chromosomal aneuploidies, based on the current clinical recommendations of CMA evaluation in ASD.

## Methods

### Study Design

Data from the 12-week multicenter, randomized pragmatic clinical trial of the Swedish version of SSGT “KONTAKT” (hereafter referred to SSGT)^12^ and genetic screening were used for this study. The initial trial tested the effectiveness of SSGT as a complement to standard care in 13 child and adolescent psychiatry outpatient units in Sweden. Outcome measures were collected at pre-, post-intervention (12-weeks) and follow-up (3-months after post-intervention). The protocol and results from the trial have been published previously.^10,12^ The study was approved by the ethical review board in Stockholm, Sweden (Dnr 2012/385-31/4) and the clinical authorities of the two involved counties. The trial was also registered online (NCT01854346). Written consent from the parents or legal guardians and verbal assent from the youth were collected. The trial and the collection of saliva samples were done between August 2012 and October 2015.

### Participants

The trial included 296 youths aged 8 to 17 years with ASD. Eligibility criteria and recruitment of participants are described in detail elsewhere.^10^ In short, participants had a clinical consensus diagnosis of autism, atypical autism, Asperger syndrome, or pervasive developmental disorder not otherwise specified based on ICD-10 criteria.^22^ The diagnosis was corroborated by ASD cutoffs (modules 3 or 4) on the Autism Diagnostic Observation Schedule (ADOS).^23^ Additionally, the ADOS total scores were used to estimate the autism severity at baseline. All participants had full-scale IQs > 70, according to the Wechsler Intelligence Scale for Children (third or fourth edition)^24^ and at least one common comorbid psychiatric ICD-10 diagnosis of ADHD, anxiety disorder, or mood disorder. Participants providing saliva samples and primary outcome data for at least one-time point (post-intervention or 3-months follow-up) were included in the planned subgroup analyzes of CNV carriers (N=209; 106 from the SSGT group, 103 from the ‘standard care’ group). We also used the baseline measures from the participants without saliva samples (n= 87) to analyze if any differences were observed between the included and excluded participants.

### Outcome Measures

The primary outcome measure was the change in total raw scores on the parent reported Social Responsiveness Scale (SRS).^25^ Higher values on the SRS scale indicate greater severity of autistic symptoms. Secondary outcome measures used in the RCT were parent ratings on the Adaptive Behavior Assessment System II (ABAS-II)^26^, the trainer-rated Developmental Disabilities modification of the Children’s Global Assessment Scale (DD-CGAS)^27^ and Ohio State University (OSU) Global Severity Scale for Autism (OSU Autism-CGI-S).^28^

### Genotyping and CNV Calling

The participants collected saliva samples using the Oragene·DNA OG-500 tubes (DNA Genotek, Inc., Ottawa, Ontario, Canada) at home using a recommended procedure. Thereafter, the DNA was extracted using Chemagen kit (PerkinElmer chemagen, Baesweiler, Germany) with chemagicSTAR^®^-robot (Hamilton Robotics, Reno, NV, USA). The genotyping of the DNA samples were done on the Affymetrix CytoScan^™^ HD microarray platform (Santa Clara, CA, USA), which has approximately 2.7 million probes, following the manufacturer’s recommendations. The identification of CNVs was made by incorporating calls from two algorithms Chromosome Analysis Suite (ChAS) software v.3.1 (Affymetrix), and Partek Genomics Suite software, version 6.6 (Partek Inc., St.Louis, MO, USA). Standard quality control was performed for the single nucleotide polymorphism (SNP) and CNV data.

For the statistical analyses, we considered only variants that were called by both algorithms and spanned at least 25 kb and 25 consecutive probes.^29^ If large CNVs were found to affect the sex chromosomes, the calls were manually inspected using ChAS software (Affymetrix) to verify the call and check for possible chromosomal aneuploidies. As we sought to limit analyses to CNVs that would be included in clinical CMA reports, we removed variants that were present with more than 0.1% frequency in the general population using the Ontario Population Genomics Platform (OPGP) data as reference.^29^ In OPGP, 873 samples were genotyped using the same Affymetrix platform. Additionally, we removed variants that overlapped more than five variants that are reported in the Database of Genomic Variants (DGV).^30^ The CNVs were mapped using GRCh37/hg19 coordinates. Thereafter, CNVs that overlapped at least one coding exon based on the gene annotation from RefSeq were included in the final CNV set. We performed experimental validation of 15 selected CNVs using quantitative polymerase chain reaction (qPCR).

### Statistical Analyses

CNV status was coded binary with “0” for non-carriers and “1” for carriers. The rare genic CNVs were categorized based on their size into three groups: 25-100 kb (small), 101-500 kb (middle-size), and >500kb including chromosome aneuploidies (large). The participants were grouped according to the largest CNV present. The effects of all rare genic CNVs, and the three CNV size groups, on the outcome were tested separately using mixed linear models (MLM). Based on our primary hypothesis that rare CNVs would affect the outcome of SSGT, we first tested the twoway interaction of CNV carrier*time (post-intervention and 3-months follow-up) in the active SSGT participant group. Thereafter, we examined if significant results would remain after including the ‘standard care’ group in a three-way interaction CNV carrier*time*intervention group (SSGT vs. ‘standard care’). Our primary MLM model included the fixed effects of age group (children or adolescents) and sex, as these had shown to influence intervention outcome in the RCT study.^10^ To account for the putative differences between participating centers and individuals, these were included as random effects in the models. The least-square (LS) means, and 95% confidence interval (CI) were calculated. The secondary model included IQ as a covariate based on previous studies suggesting an association of CNVs with IQ and educational attainment in the general population.^31,32^ The effect sizes were calculated by dividing the group difference in the change of least-squares means from pre-intervention to post-intervention/follow-up by the pooled standard deviation at pre-intervention. Additionally, we tested for differences in clinical group characteristics at baseline (SRS, IQ, ADOS total score) between the participants included and excluded in this study within the SSGT and standard care group as well as between rare genic CNV carriers and non-carriers using two-sided student’s t-test and χ^2^ test. The analyses were conducted using R version 3.4.2.

## Results

Demographics and clinical characteristics of the participants included in this study are presented in Table 1, stratified by intervention group. After genotyping and CNV quality control two samples (one from SSGT and one from standard care group) were removed. Therefore, 105 children were included in the active SSGT group and 102 in the standard care group. Eighty-seven participants (29.3%) from the RCT were excluded due to unavailable saliva sample for the genetic analysis. IQ was significantly lower in the SSGT participants without a saliva sample (eTable 1). No other differences in the baseline measures were seen between the included and excluded participants (eTable 1). The participants without saliva samples had a higher frequency of missing primary outcome data in both SSGT and standard care groups (eTable 2).

**Table 1.** Demographic and Clinical Characteristics at Baseline by Intervention Group.

| Characteristics | SSGT (n=105) | Standard care (n=102) |
| --- | --- | --- |
| Sex |  |  |
| Female, n (%) | 28 (26.4%) | 32 (31.4%) |
| Male, n (%) | 77 (73.3%) | 70 (68.6%) |
| Age (mean, SD) | 11.92 ($\pm 2.59$) | 11.48 ($\pm 2.69$) |
| Adolescents, n (%) | 39 (37.1%) | 37 (36.3%) |
| Children, n (%) | 66 (62.9%) | 65 (63.7%) |
| Comorbidity |  |  |
| ADHD, n (%) | 77 (73.0%) | 75 (73.5%) |
| Anxiety, n (%) | 23 (22.0%) | 23 (22.5%) |
| Depression, n (%) | 13 (12.4%) | 16 (15.7%) |
| Others, n (%) | 16 (15.2%) | 14 (13.7%) |
| 1 Comorbidity, n (%) | 83 (79.0%) | 77 (75.5%) |
| 2 or more comorbidities, n (%) | 22 (21.0%) | 25 (24.5%) |
| Cognitive and Behavioral Measures |  |  |
| IQ (mean, SD) | 98.48 ($\pm 13.08$) | 98.43 ($\pm 12.94$) |
| SRS pre-intervention (mean, SD) | 86.08 ($\pm 25.06$) | 86.33 ($\pm 24.30$) |
| ADOS total score (mean, SD) | 10.60 ($\pm 3.31$) | 10.95 ($\pm 3.45$) |
| DD-CGAS (mean, SD) | 58.26 ($\pm 7.27$) | 57.13 ($\pm 7.02$) |
| OSU Autism CGI-S (mean, SD) | 4.23 ($\pm 0.83$) | 4.41 ($\pm 0.79$) |
| ABAS-II (mean, SD) | 371.95 ($\pm 73.0$) | 374.66 ($\pm 61.89$) |

Abbreviations: SSGT, Social Skills Group Training; ADHD, attention-deficit hyperactivity disorder; SRS, Social Responsiveness Scale; ADOS, the Autism Diagnostic Observation Schedule; DD-CGAS, Developmental Disabilities modification of the Children's Global Assessment Scale; OSU Autism CGI-S, Ohio State University (OSU) Global Severity Scale for Autism; ABAS-II the Adaptive Behavior Assessment System II.

Of the 207 included participants, 71 (34.8%) carried at least one rare genic CNV larger than 25 kb. Of these, 14 (6.8%) were carriers of large CNVs including the chromosomal aneuploidies, 42 (20.3%) carriers of middle-size CNVs and 15 (7.2%) carriers of small CNVs.

Three participants (1.45%) were carriers of sex chromosome aneuploidies (47,XXX; 45,X; 47;XYY). Experimental validation using qPCR was done for 18% of the identified rare variants with 100% validation rate. The CNV carriers in the SSGT group had significantly lower average IQ in comparison with the non-carriers (T = 2.02, *P*= .047) but not in the whole sample (T = 1.67, p = .097) (eFigure 1). No differences at baseline were found for SRS total raw scores or ADOS total scores between the CNV carriers and non-carriers (eFigure 1). The majority of rare genic CNVs identified in the participants would be reported from CMA to the participants in clinical settings including 15 clinically significant variants (15/207, 7.2%). These include the three sex chromosome syndromes, large deletion and duplications affecting known risk CNV loci (15q11.2-q13.1; 9p24.3-p23; 7q11.2; 15q26.2-q26.3; Xq27.3-Xq27.2), and smaller CNVs affecting genes with strong evidence to be involved in ASD or related conditions such as *GATAD2B* (OMIM *614998), *CHD8* (OMIM*610528), *ASH1L* (OMIM*607999), *KDM6A* (OMIM*300128), and *KDM5B*(OMIM*605393).

The sex and age group (children or adolescents) adjusted results for the primary outcome are reported in Table 2, and the estimated LS-means are shown in Figure 1. No differences were found for the carriers of rare genic CNVs at post-intervention or follow-up in comparison with non-carriers. We then stratified the data by size of the CNVs, the carriers of large genic CNVs (>500 kb), including chromosomal aneuploidies, had inferior outcome at post-intervention (β = 15.35, 95% CI 2.9-27.8, *P* = .017) and at follow-up (β = 14.19, 95% CI 1.7-26.7, *P* = .028) compared with participants without large genic CNVs (Table 2, Figure 1). Changes between pre- and post-intervention/follow-up for the carriers and non-carries are shown in eFigure 2. We observed medium effect sizes for both time points (−0.47 at post and −0.43 at follow-up) showing the negative impact of large CNVs on the outcome in comparison with participants without large CNVs. An opposite effect, however, was observed for carriers of middle-size CNVs at follow-up (β =-8.69, 95% CI −16.8 - −0.6, *P* = .038) with an effect size of 0.28. In the three-way interaction model, when including the standard care group to assure the specificity of effect for SSGT, the significant effect for large genic CNV carriers in SSGT group remained at post-intervention (β = 21.90, 95% CI 1.5 - 42.3, *P* = .036) with a similar trend at follow-up (β =18.82, 95% CI −1.6 – 39.3, *P* = .072) (Table 2, Figure 1). The model adjusted for IQ showed similar results (eTable 3).

**Table 2.** Model Estimates of Associations Between Rare Copy Number Variation and Primary Outcome.

|  | SSGT (n=105) |  |  | Total Sample (N=207) |  |  |
| --- | --- | --- | --- | --- | --- | --- |
| Mixed Linear Model | $\beta$ (95% CI) | P value | Effect Size (95% CI)<sup>a</sup> | $\beta$ (95% CI) | P value | Effect Size (95% CI)<sup>a</sup> |
| Carrier of a rare CNV |  |  |  |  |  |  |
| *SRSpst | 5.27 (-2.1, 12.6) | .16 | - | -1.66 (-9.0, 5.7) | .66 | - |
| *SRSfu | -2.59 (-10.0, 4.9) | .50 | - | -0.02 (-7.8, 7.8) | 1.0 | - |
| *SRSpst*SSGT | - | - | - | 6.99 (-3.1- 17.1) | .18 | - |
| *SRSfu*SSGT | - | - | - | -2.52 (-13.0, 8.0) | .64 | - |
| Carrier of a rare large CNV (>500 kb) |  |  |  |  |  |  |
| *SRSpst | 15.35 (2.9, 27.8) | .017 | -0.47 (-1.12, 0.19) | -6.7 (-23.3, 10.0) | .43 | - |
| *SRSfu | 14.19 (1.7, 26.7) | .028 | -0.43 (-1.09, 0.22) | -4.57 (-21.24, 12.1) | .59 | - |
| *SRSpst*SSGT | - | - | - | 21.90 (1.5, 42.3) | .036 | -0.55 (-1.37, -0.06) |
| *SRSfu*SSGT | - | - | - | 18.82 (-1.6, 39.3) | .072 | - |
| Carrier of middle size CNV (100-500 kb) |  |  |  |  |  |  |
| *SRSpst | -0.77 (-8.8, 7.3) | .85 | - | -2.05 (-11.2, 7.1) | .66 | - |
| *SRSfu | -8.69 (-16.8, -0.6) | .038 | 0.28 (-0.18, 0.74) | -3.46 (-12.9, 6.0) | .47 | - |
| *SRSpst*SSGT | . | - | - | 1.32 (-10.6, 13.2) | .83 | - |
| *SRSfu*SSGT | - | - | - | -5.25 (-17.4, 6.9) | .40 | - |
| Carrier of small CNV (20-100 kb) |  |  |  |  |  |  |
| *SRSpst | 2.93 (-13.1, 18.9) | .72 | - | 2.32 (-9.2, 13.8) | .69 | - |
| *SRSfu | - 0.78 (-20.1, 18.6) | .94 | - | 11.00 (-3.2, 25.2) | .13 | - |
| *SRSpst*SSGT | - | - | - | 0.93 (-17.9, 19.8) | .92 | - |
| *SRSfu*SSGT | - | - | - | -11.58 (-34.6, 11.4) | .33 | - |

Abbreviations: SRSpst, Social Responsiveness scale outcome at post-intervention; SRSfu, Social Responsiveness Scale outcome at follow-up;SSGT, Social skills group training. “-” indicates not calculated. <sup>a</sup> Effect sizes were calculated only for the significant models.

**Figure 1.**
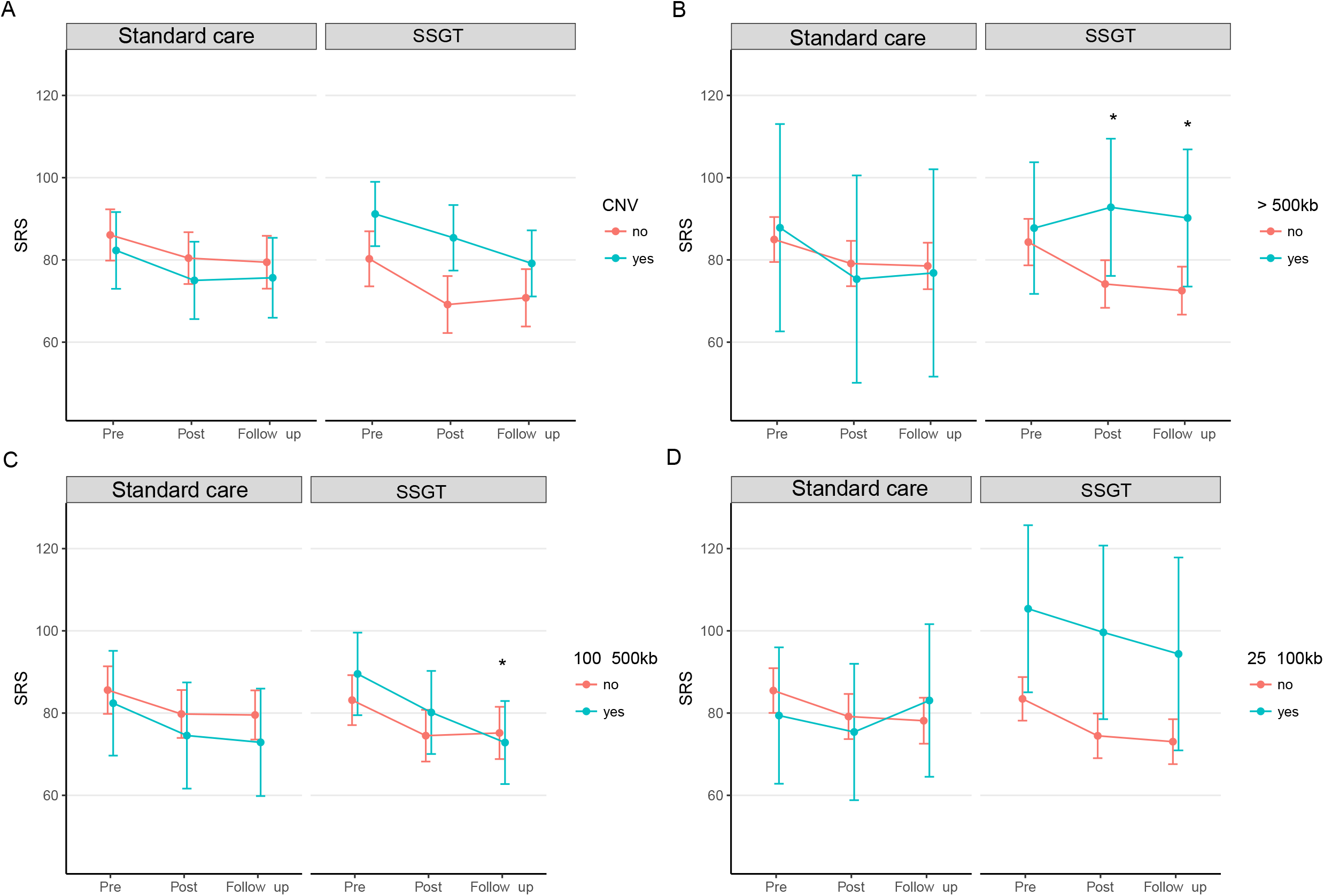
Least-square means and 95% confidence interval for mixed-linear model for parent-rated Social Responsiveness Scale (SRS) scores for the standard care and SSGT groups stratified to carriers of rare genic copy number variation (CNV) and non-carriers (A) and based on the size of the CNV; >500 kb and chromosomal aneuploidies (B), 101 - 500 kb (C) and 25 kb - 100 kb (D).

The results for the secondary outcomes, parent-rated ABAS-II, trainer-rated DD-CGAS and OSU Autism CGI-S, are presented in eTable 4. Similar to SRS results, carriers of large rare genic CNVs showed inferior outcome as measured by ABAS-II at post-intervention (β = −28.75, 95 % CI −56.6 - −0.9 *P* = .045), however not remaining significant in the three-way interaction model (β = −40.39, 95 % CI −84.5 – 3.7 *P* = .073). For the trainer-ratings using OSU Autism CGI S, carriers of small CNVs showed greater improvements at post-intervention (β = −1.20, 95 % CI −2.0 - −0.43 *P* = .003) with significant three-way interaction model at same time point (β = −1.21, 95 % CI −2.1 - −0.3 *P* = .008). The IQ adjusted model showed similar results (eTable 5).

## Discussion

This is the first study to analyze the moderating effects of rare genic CNVs on SSGT intervention in children and adolescents with ASD. Our results suggest that in general being a carrier of any size rare genic CNV did not impact the primary outcome. However, the prespecified subgroup analyzes showed that carriers of large genic CNVs (>500 kb), including chromosome aneuploidies, improve less than non-carriers. Previous studies have reported that many of the recurrent large rare CNVs are associated with low IQ,^31,32^ but our adjusted models indicated effects of CNVs above and beyond IQ. The inferior outcome of carriers of large rare genic CNVs was robust when analyzing parent-rated primary and secondary outcome measures but were not replicated in trainer-rated secondary outcomes. The results indicate that this type of information eventually might help guide personalized intervention planning and clinical resource allocation.

The recent molecular evidence in ASD hopefully can be used to design effective medical interventions targeting ASD core symptoms.^33^ However, there is still a long road ahead until such interventions can be introduced on the market and implemented in clinical practice. Therefore, it is crucial to investigate how genetic information can help to enhance treatment decisions using current and widely applied behavioral ASD programs. Studies on potential genetic moderators of both pharmacological (pharmacogenomics) and psychological treatments (therapygenetics) in psychiatry has started to emerge in recent years.^34,35^ For instance, a recent study showed that individuals with bipolar disorder and high genetic load as measured by polygenic scores for schizophrenia had an inadequate response to lithium treatment.^36^ Additionally, a study investigating the association of CNVs with response to antidepressant medication in major depressive disorder showed no general effect, but the authors noted less response for carriers of CNVs in specific loci, including *NRXN1*.^37^ For therapygenetics, recent studies have mostly used common genetic variants either in selected candidate genes^38^ or a hypothesis-free genome-wide approach^39^ with no or limited evidence of genetic moderators of treatment response. We used a more refined approach, by investigating rare genic CNVs that are associated with a large risk for ASD as well as other neurodevelopmental and psychiatric disorders.^40,41^ To the best of our knowledge, our study is the first to investigate therapygenetics in ASD.

Our data also show that CMA is useful in testing for genetic alterations in autistic individuals in the average and high intellectual range. We identified rates of clinically significant CNVs (7.2%) similar to what has previously been reported in samples including individuals with ASD in the intellectual disability range.^17-19^ Interestingly, our results seem to corroborate previous studies showing that individuals with ASD and genomic or monogenic syndromes form a subgroup characterized by lower cognitive abilities and milder ASD symptomatology, especially for social communication alterations.^42,43^ Similarly, an earlier study using machine learning classification showed that individuals with specified genetic disorders in ASD have a specific signature of social impairment.^44^ Therefore, it is crucial to better understand the intervention needs of those with specified molecular alterations. For instance, it should not be ruled out that this group could benefit from longer periods of training. We recently reported that a long version of KONTAKT (24-weeks) resulted in larger effects in general than we have found for the 12-weeks version used here.^45^ Unfortunately, that trial was not sufficiently powered for subgroup analyzes.

Limitations of our study should be noted. First, while this study was based on the largest RCT in ASD research to date, the sample was still limited to investigate effects of single loci. Thus, our results reflect various molecular causes. Second, data on either saliva or the primary outcome were missing for 87 (29%) of the participants enrolled in the trial, and these participants from the active SSGT group had significantly lower IQ. Third, the secondary outcome measures provided a somewhat inconsistent picture, underscoring the preliminary nature of the results. Fourth, the showed subgroup differences should be interpreted with caution. These limitations must be addressed in future studies before specific clinical recommendations can be made based on CNVs. From a clinical and practical perspective, we would also like to stress that there are many social, economical, legal and ethical issues to be addressed as the genetic and genomic testing in ASD is rapidly increasing. These include, but are not limited to, realistic cost-benefit analyses, genetic data protection, clinical guideline development, and the avoidance of discrimination of genetic and ethnic minorities.^46,47,6^

## Conclusions

This is the first study to show that individuals with ASD and large rare genic CNVs are less likely than non-carriers to benefit from SSGT. Thus, a combination of CMA results and other demographic and clinical information (e.g., age, sex, IQ, comorbidity, social communication severity) might have the potential to determine the likelihood of none, or low, response. Until replicated in independent samples, our results should be interpreted with caution.

## Data sharing

The raw array data or individual CNV calls have not been shared in public database due to limited ethical approval and informed consent from the participants for data sharing but are available from the corresponding author upon reasonable request and subject to necessary clearances.

## Author Contributions

*Study Concept and Design*: Tammimies, Choque-Olsson, and Bölte

*Acquisition, Analysis, or Interpretation of Data*: All authors

*Drafting Manuscript*: Tammimies, Li, and Rabkina

*Critical Revision of the Manuscript for Important Intellectual Content:* All authors

*Statistical Analysis*: Tammimies, Li, Rabkina

*Obtained Funding*: Tammimies, Bölte

*Administrative, Technical, or Material support*, Rabkina, Stamouli, Becker, Nicolaou, Berggren, Coco, Falkmer, Jonsson, Choque-Olsson

*Study Supervision*: Tammimies, Bölte

## Acknowledgments

We thank the children, adolescents, and parents who participated in the study. Oskar Flygare, MSc, Anders Görling MSc and Kerstin Andersson MSc are acknowledged for their work in collecting the data and samples during the RCT. The authors are also thankful to the leads of child and adolescent psychiatry (Olav Bengtsson, MD, Paula Liljeberg, MD, Charlotta Wiberg Spangenberg, MSc, Peter Ericson, MSc, Karin Forler, MSc, Alkisti Nikolayidis Linderholm, MSc, all of Stockholm County Council), PRIMA Järva child and adolescent psychiatry (MaiBritt Giacobini, MD, PhD), and child and adolescent habilitation services (Lars Kjellin, PhD, Moa Pellrud, MSc, of Örebro County Council) for organizational support. The genotyping was done at Aros Applied Biotechnology in Denmark. The computation resources were provided by SNIC through Uppsala Multidisciplinary Center for Advanced Computational Science (UPPMAX).

## References

1. Idring S, Lundberg M, Sturm H, et al. Changes in prevalence of autism spectrum disorders in 2001-2011: findings from the Stockholm youth cohort. Journal of autism and developmental disorders. 2015;45(6):1766–1773.

2. Identified prevalence of autism spectrum disorder. Centers for Disease Control and Prevention (CDC) 2018. Accessed 30 May 2018.

3. Diagnostic and statistical manual of mental disorders. In. Vol 5th ed: American Psychiatric Association; 2013.

4. Simonoff E, Pickles A, Charman T, Chandler S, Loucas T, Baird G. Psychiatric disorders in children with autism spectrum disorders: prevalence, comorbidity, and associated factors in a population-derived sample. J Am Acad Child Adolesc Psychiatry. 2008;47(8):921–929.

5. Alexeeff SE, Yau V, Qian Y, et al. Medical Conditions in the First Years of Life Associated with Future Diagnosis of ASD in Children. Journal of autism and developmental disorders. 2017;47(7):2067–2079.

6. Woodbury-Smith M, Scherer SW. Progress in the genetics of autism spectrum disorder. Developmental medicine and child neurology. 2018;60(5):445–451.

7. Hertz-Picciotto I, Schmidt RJ, Krakowiak P. Understanding environmental contributions to autism: Causal concepts and the state of science. Autism research : official journal of the International Society for Autism Research. 2018;11(4):554–586.

8. Wong C, Odom SL, Hume KA, et al. Evidence-Based Practices for Children, Youth, and Young Adults with Autism Spectrum Disorder: A Comprehensive Review. Journal of autism and developmental disorders. 2015;45(7):1951–1966.

9. Jonsson U, Choque Olsson N, Bölte S. Can findings from randomized controlled trials of social skills training in autism spectrum disorder be generalized? The neglected dimension of external validity. Autism : the international journal of research and practice. 2016;20(3):295–305.

10. Choque Olsson N, Flygare O, Coco C, et al. Social Skills Training for Children and Adolescents With Autism Spectrum Disorder: A Randomized Controlled Trial. J Am Acad Child Adolesc Psychiatry. 2017;56(7):585–592.

11. Freitag CM, Jensen K, Elsuni L, et al. Group-based cognitive behavioural psychotherapy for children and adolescents with ASD: the randomized, multicentre, controlled SOSTA-net trial. J Child Psychol Psychiatry. 2016;57(5):596–605.

12. Choque-Olsson N, Tammimies K, Bölte S. Social skills group training for children and adolescents, “KONTAKT”, multicenter, randomized controlled trial: protocol. Translational Developmental Psychiatry. 2015;3: 29825.

13. Herbrecht E, Poustka F, Birnkammer S, et al. Pilot evaluation of the Frankfurt Social Skills Training for children and adolescents with autism spectrum disorder. European child & adolescent psychiatry. 2009;18(6):327–335.

14. Deckers A, Muris P, Roelofs J, Arntz A. A Group-Administered social Skills Training for 8-to 12-Year-Old, high-Functioning Children With Autism Spectrum Disorders: An Evaluation of its Effectiveness in a Naturalistic Outpatient Treatment Setting. Journal of autism and developmental disorders. 2016;46(11):3493–3504.

15. Pinto D, Delaby E, Merico D, et al. Convergence of genes and cellular pathways dysregulated in autism spectrum disorders. American journal of human genetics. 2014;94(5):677–694.

16. Yuen R, Merico D, Bookman M, et al. Whole genome sequencing resource identifies 18 new candidate genes for autism spectrum disorder. Nature neuroscience. 2017;20(4):602–611.

17. Tammimies K, Marshall CR, Walker S, et al. Molecular Diagnostic Yield of Chromosomal Microarray Analysis and Whole-Exome Sequencing in Children With Autism Spectrum Disorder. Jama. 2015;314(9):895–903.

18. Ho KS, Wassman ER, Baxter AL, et al. Chromosomal Microarray Analysis of Consecutive Individuals with Autism Spectrum Disorders Using an Ultra-High Resolution Chromosomal Microarray Optimized for Neurodevelopmental Disorders. International journal of molecular sciences. 2016;17(12).

19. Roberts JL, Hovanes K, Dasouki M, Manzardo AM, Butler MG. Chromosomal microarray analysis of consecutive individuals with autism spectrum disorders or learning disability presenting for genetic services. Gene. 2014;535(1):70–78.

20. Hayeems RZ, Hoang N, Chenier S, et al. Capturing the clinical utility of genomic testing: medical recommendations following pediatric microarray. European journal of human genetics : EJHG. 2015;23(9):1135–1141.

21. Henderson LB, Applegate CD, Wohler E, Sheridan MB, Hoover-Fong J, Batista DA. The impact of chromosomal microarray on clinical management: a retrospective analysis. Genetics in medicine : official journal of the American College of Medical Genetics. 2014;16(9):657–664.

22. World Health Organization. International statistical classification of diseases and related health problems, 10th Revision (ICD-10). Geneva1992.

23. Lord C, Risi S, Lambrecht L, et al. The autism diagnostic observation schedule-generic: a standard measure of social and communication deficits associated with the spectrum of autism. Journal of autism and developmental disorders. 2000;30(3):205–223.

24. Wechsler D, Kaplan E, Fein D, et al. Wechsler Intelligence Scale for Children-Fourth Edition-Integrated. San Antonio, TX: Harcourt. 2004.

25. Constantino JN, Gruber CP. Social Responsiveness Scale (SRS). Vol Los Angeles CA: Western Psychological Services; 2005.

26. Harrison PL, Oakland T. Adaptive behavior assessment system (2nd ed.). San Antonio, : TX: Harcourt Assessment, Inc.; 2003.

27. Wagner A, Lecavalier L, Arnold LE, et al. Developmental disabilities modification of the Children’s Global Assessment Scale. Biological psychiatry. 2007;61(4):504–511.

28. OSU Research Unit on Pediatric Psychopharmacology. OSU Autism Rating Scale CGI-DSM-IV (OARS-4). OSU Autism Rating Scale-CGI-psychmed. 2005.

29. Uddin M, Thiruvahindrapuram B, Walker S, et al. A high-resolution copy-number variation resource for clinical and population genetics. Genetics in medicine : official journal of the American College of Medical Genetics. 2015;17(9):747–752.

30. MacDonald JR, Ziman R, Yuen RK, Feuk L, Scherer SW. The Database of Genomic Variants: a curated collection of structural variation in the human genome. Nucleic acids research. 2014;42(Database issue):D986–992.

31. Merikangas AK, Segurado R, Heron EA, et al. The phenotypic manifestations of rare genic CNVs in autism spectrum disorder. Molecular psychiatry. 2015;20(11):1366–1372.

32. Mannik K, Magi R, Mace A, et al. Copy number variations and cognitive phenotypes in unselected populations. Jama. 2015;313(20):2044–2054.

33. Loth E, Spooren W, Ham LM, et al. Identification and validation of biomarkers for autism spectrum disorders. Nat Rev Drug Discov. 2016;15(1):70–73.

34. Pisanu C, Heilbronner U, Squassina A. The Role of Pharmacogenomics in Bipolar Disorder: Moving Towards Precision Medicine. Mol Diagn Ther. 2018.

35. Eley TC. The future of therapygenetics: where will studies predicting psychological treatment response from genomic markers lead? Depress Anxiety. 2014;31(8):617–620.

36. International Consortium on Lithium Genetics, Amare AT, Schubert KO, et al. Association of Polygenic Score for Schizophrenia and HLA Antigen and Inflammation Genes With Response to Lithium in Bipolar Affective Disorder: A Genome-Wide Association Study. JAMA psychiatry. 2018;75(1):65–74.

37. Tansey KE, Rucker JJ, Kavanagh DH, et al. Copy number variants and therapeutic response to antidepressant medication in major depressive disorder. Pharmacogenomics J. 2014;14(4):395–399.

38. Lester KJ, Coleman JR, Roberts S, et al. Genetic variation in the endocannabinoid system and response to Cognitive Behavior Therapy for child anxiety disorders. American journal of medical genetics Part B, Neuropsychiatric genetics : the official publication of the International Society of Psychiatric Genetics. 2017;174(2):144–155.

39. Coleman JR, Lester KJ, Keers R, et al. Genome-wide association study of response to cognitive-behavioural therapy in children with anxiety disorders. The British journal of psychiatry : the journal of mental science. 2016;209(3):236–243.

40. Muhle RA, Reed HE, Stratigos KA, Veenstra-VanderWeele J. The Emerging Clinical Neuroscience of Autism Spectrum Disorder: A Review. JAMA psychiatry. 2018;75(5):514–523.

41. Lowther C, Costain G, Baribeau DA, Bassett AS. Genomic Disorders in Psychiatry-What Does the Clinician Need to Know? Curr Psychiatry Rep. 2017;19(11):82.

42. Bishop SL, Farmer C, Bal V, et al. Identification of Developmental and Behavioral Markers Associated With Genetic Abnormalities in Autism Spectrum Disorder. The American journal of psychiatry. 2017;174(6):576–585.

43. Iossifov I, O’Roak BJ, Sanders SJ, et al. The contribution of de novo coding mutations to autism spectrum disorder. Nature. 2014;515(7526):216–221.

44. Bruining H, Eijkemans MJ, Kas MJ, Curran SR, Vorstman JA, Bolton PF. Behavioral signatures related to genetic disorders in autism. Molecular autism. 2014;5(1):11.

45. Jonsson U, Olsson NC, Coco C, et al. Long-term social skills group training for children and adolescents with autism spectrum disorder: a randomized controlled trial. European child & adolescent psychiatry. 2018.

46. Botkin JR, Belmont JW, Berg JS, et al. Points to Consider: Ethical, Legal, and Psychosocial Implications of Genetic Testing in Children and Adolescents. American journal of human genetics. 2015;97(1):6–21.

47. Yuen T, Carter MT, Szatmari P, Ungar WJ. Cost-effectiveness of Genome and Exome Sequencing in Children Diagnosed with Autism Spectrum Disorder. Appl Health Econ Health Policy. 2018;16(4):481–493.

